# Heterogeneous hybrid immunity against Omicron variant JN.1 at 11 months following breakthrough infection

**DOI:** 10.1101/2024.03.02.583082

**Authors:** Xuan He, Jiajing Jiang, Guo Li, Jinyuan Liu, Jiadi Gan, Linlin Zhou, Chunyang Bai, Qiong Zi, Xiaoli Mou, Shan Zeng, Junjie Yuan, Chuanjie Zhou, Yangqian Li, Guonian Zhu, Renjie Zhao, Lan Yang, Jiaxuan Wu, Huohuo Zhang, Jinghong Xian, Zhoufeng Wang, Qi Qi, Yu Liu, Jingyou Yu, Dan Liu, Weimin Li

## Abstract

A highly transmissible SARS-CoV-2 variant JN.1 is rapidly spreading throughout the nation, becoming the predominant strain in China and worldwide. However, the current immunity against the circulating JN.1 at population level has yet to be fully evaluated. We recruited representative cohorts with stratified age groups and diverse combinations of vaccination and/or infection in recent months, and promptly assessed humoral immunity for these subjects predominantly exhibiting hybrid immunity. We report that at 11 months following BA.5-wave breakthrough infection (BTI), these vaccinated individuals generally showed above-the-threshold yet low level of neutralizing activity against JN.1, with slightly greater potency observed in children and adolescents compared to adults and seniors. Meanwhile, XBB/EG.5-wave reinfection post-BTI significantly boosted the neutralizing antibodies against Omicron variants, including JN.1 in both adults (13.4-fold increase) and seniors (24.9-fold increase). To better understand respiratory mucosal protection against JN.1 over an extended period of months post-BTI, we profiled the humoral immunity in bronchoalveolar lavage samples obtained from vaccinated subjects with or without BTI, and revealed increased potency of neutralizing activity against the BA.5 and JN.1 variants in the respiratory mucosa through natural infection. Notably, at 11 months post-BTI, memory B cell responses against prototype and JN.1 were detectable in both blood and respiratory mucosa, displaying distinct memory features in the circulation and airway compartments. XBB/EG.5-wave reinfection drove the expansion of JN.1-specific B cells, along with the back-boosting of B cells responding to the ancestral viral strain, suggesting the involvement of immune imprinting. Together, this study indicates heterogeneous hybrid immunity over 11 months post-BTI, and underscores the vulnerability of individuals, particularly high-risk seniors, to JN.1 breakthrough infection. An additional booster with XBB-containing vaccine may greatly alleviate the onward transmission of immune-evasive SARS-CoV-2 variants.

## Introduction

Coronavirus disease-2019 (COVID-19) is a novel respiratory illness caused by a highly pathogenic and transmissible severe acute respiratory syndrome-coronavirus-2 (SARS-CoV-2), leading to widespread illness and death globally.^1^ While the World Health Organization (WHO) has officially declared the end of the COVID-19 pandemic, SARS-CoV-2 persists, spreading and mutating within human populations. Since the emergence of the Omicron (B.1.1.529) which was designated as variant of concern (VOC) by WHO in November 2021, the Omicron lineages have swiftly traversed the globe, continuously circulating, evolving and giving rise to subvariants. Although Omicron variants generally exhibit lower pathogenicity as compared with other SARS-CoV-2 lineages, they demonstrate enhanced transmissibility and a higher capacity of immune escape.^2^ The subvariant of Omicron (BA.2.86) was initially identified in August 2023, distinguishing itself from other prevalent variants with over 30 additional mutations in the spike protein, including 15 mutations located in the receptor-binding domain (RBD).^3^ Previous studies have demonstrated that BA.2.86 is one of the most rapidly spreading and immune evasive mutants.^4, 5^ Recently, a sub-lineage of the BA.2.86, known as JN.1, has rapidly surged within a few weeks, promoting the WHO to step up its classification as variant of interest (VOI) in mid-December.^6^ L455S, a mutation prominently associated with immune evasion concerns, is regarded as the one of the most characteristic mutations in JN.1 variant as compared with its predecessor BA.2.86. By compensating the weakness of BA.2.86 against class 1 neutralizing antibodies via the additional acquisition of L455S, JN.1 significantly enhances its resistance to humoral immune response against SARS-CoV-2, surpassing both BA.2.86 and EG.5 lineages.^4, 7^ Besides, the pseudovirus assay indicated the superior infectivity of JN.1 compared to BA.2.86, although there is no solid evidence that JN.1 causes more severe disease.^7^ Data from the Centers for Disease Control and Prevention (CDC) in the United States indicate that since late December 2023, JN.1 has quickly overtaken other Omicron variants as the dominant strain in the country. ^8^

Vaccination stands as one of the primary methods for containing viral infections and preventing the spread of infectious diseases. However, challenges such as immune evasion and waning immunity, as seen in the case of SARS-CoV-2 infection, undermine our immunization endeavors. Consequently, individuals face an increasing risk of breakthrough infections, being associated with the development of hybrid immunity, an immunity induced by combination of vaccination and natural infection.^9, 10^ Furthermore, mismatches between the vaccine regimen and infection strain may further complicate the efficacy of immune protection, likely due to the phenomenon known as immune imprinting. After BA.5 infection, regardless of the vaccination status, the neutralizing capacity of convalescent sera against newly emerging XBB and its subvariants is weak.^11^ However, repeated exposures to Omicron, on the other hand, have been demonstrated to markedly enhance neutralization against the Omicron variants.^12^ Moreover, immune responses may vary across groups of different ages. Following SARS-CoV-2 infection, children develop persistent antibodies and cellular immunity,^13^ but it is yet to be determined whether the humoral immunity in children is still effective against newly emerging Omicron variant JN.1, particularly over several months following vaccination or breakthrough infection. COVID-19-related morbidity and mortality are nearly absent in children, while being notably higher in the seniors.^14, 15^ As breakthrough infections have become more prevalent at population level, it is crucial to investigate the immunity against newly emerging Omicron variants in high-risk seniors.

Given the prevalence of hybrid immunity within the population, we aimed to fill gaps in the understanding of immunity against the circulating JN.1 variant in individuals with heterologous combinations of vaccination/infection and demographic profiles. This study holds significant implications for shaping future public health policies and guiding COVID-19 vaccine development.

## Results

### COVID-19 study cohorts

JN.1, which emerged from the variant BA.2.86 and was first detected in 2023 fall, currently accounted for over 80 percent of COVID-19 cases in China and the United States (Fig. 1A).^16^ JN.1 variant evolves with more than 50 substitutions on the spike region when compared with the ancestral strain, representing the most phylogenetically divergent variant as of the current stage (Fig. 1B). Despite of only one additional amino acid on position 455 (L455S) on top of BA.2.86, JN.1 showed elevated spreading capacity, which may be associated with immune escape from the vaccine-mediated protection. In line with this hypothesis, recent studies revealed enhanced immune evasion by JN.1 over its predecessor BA.2.86. ^4, 7^ Therefore, comprehensive assessment of immunity against newly emerged variant JN.1 at population level, in particular under varied COVID-19 immunization and/or infection scenarios, is warranted.

**Figure 1.**
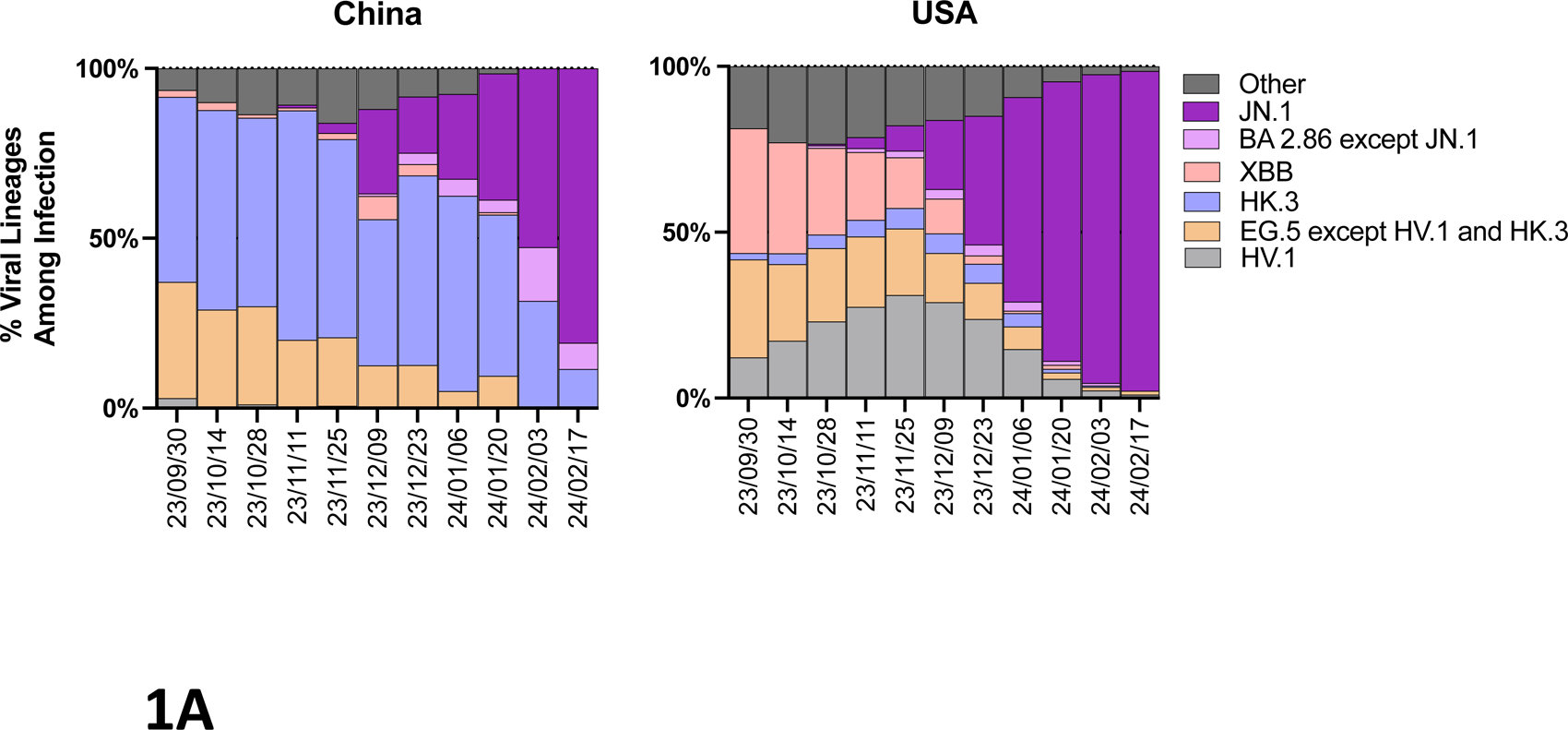
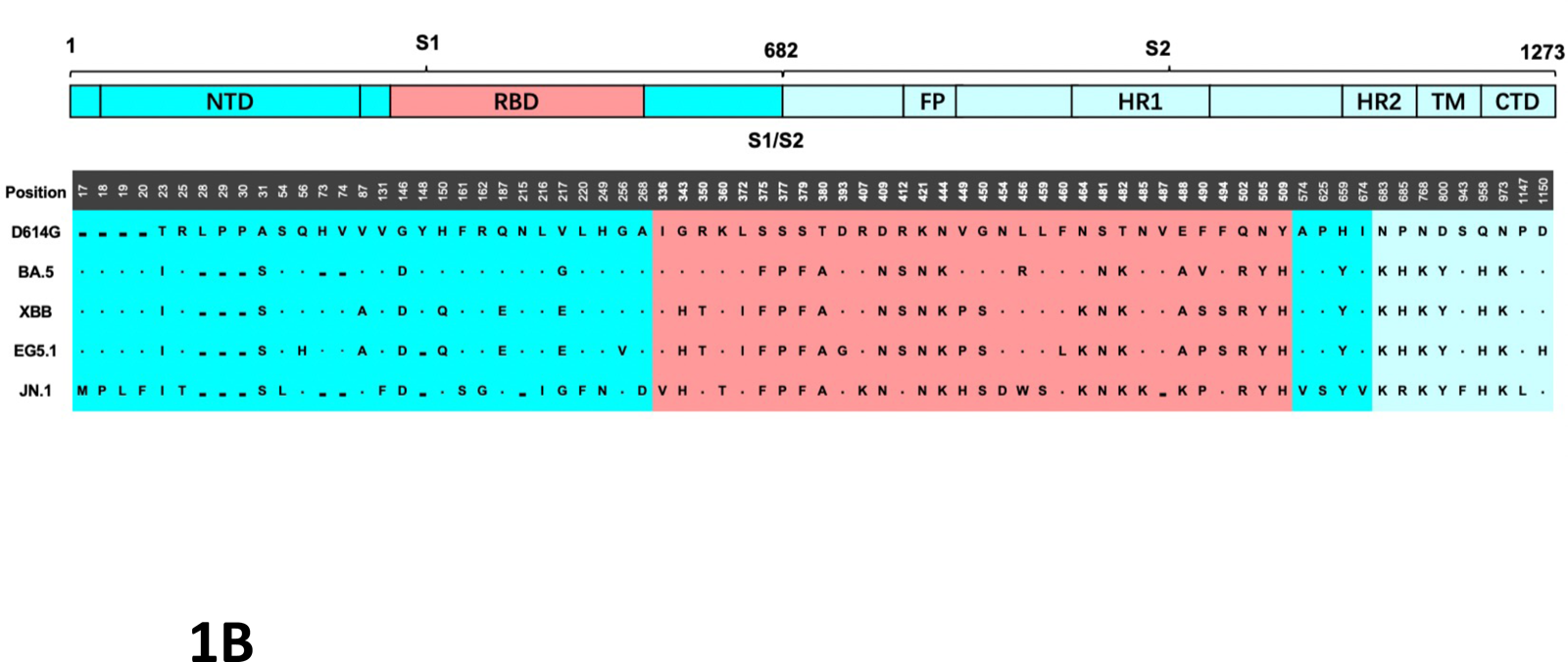
Schematic and frequency of the most abundant lineages across USA and China within the last 5 months. (**A**) The frequency of each variant is calculated as a centered 14-day rolling average. (**B**) The most representative amino acid mutations of each variant were selected to construct pseudotyped virus for this study. Each mutation in included SARS-CoV-2 variant is indicated relative to the reference D614G sequence. A dot indicates an identical amino acid at the indicated position, while a dash indicates a deletion at that point. NTD N-terminal domain, RBD Receptor-binding domain, FP Fusion peptide, HR1 Heptad repeat 1, HR2 Heptad repeat 2, TM transmembrane domain, CTD C-terminal domain.

146 individuals with COVID-19 vaccination and/or infection were recruited for this study from multiple sites throughout the Sichuan province in China (Table S1). The study primarily consisted of mild disease cases in vaccinated subjects (with at least one dose of inactivated vaccine) with breakthrough infection (BTI) during BA.5-wave occurring from December 2022 to February 2023 in Sichuan. Samples including blood and bronchoalveolar lavage (BAL) were collected between November 2023 to January 2024. For those subjects experiencing reinfection (RI) by XBB/EG.5-wave between May 2023 and June 2023 in Sichuan, blood samples were collected within two weeks post-symptom onset. These COVID-19 convalescents were grouped by age and immunization history as: unvaccinated children with BTI (n=4); vaccinated children with BTI (n=10); vaccinated adolescents with BTI (n=14); vaccinated adults with BTI (n=30); vaccinated seniors with BTI (n=30); vaccinated adults with BTI plus RI (n=13); vaccinated seniors with BTI plus RI (n=8).

### Negligible circulating neutralizing antibodies against JN.1 variant at 11 months following BA.5-wave breakthrough infection

To determine the humoral immunity against SARS-CoV-2 variants following vaccination and/or infection, we collected blood samples from unvaccinated and vaccinated individuals with BTI. Given Spike receptor binding domain (RBD) is the pivotal target for most NAbs against SARS-CoV-2,^18, 19, 20^ we performed enzyme-linked immunosorbent assay (ELISA) to assess the SARS-CoV-2 Spike RBD-specific immunoglobulin G (IgG) endpoint antibody titers against ancestral strain and primary Omicron subvariants including BA.5, XBB, EG.5.1 and JN.1 in plasma for the cohorts. By 11 months following the COVID-19 wave provoked by Omicron BA.5 subvariants, RBD-specific binding antibody responses were observed with ELISA titers of 1619 (interquartile range [IQR]: 805 to 3042) against EG.5.1 and titers of 972 (IQR: 569 to 1751) against JN.1, both of which were significantly lower than that against prototype with titers of 3686 (IQR: 1966 to 7421) (P<0.0001 for both EG.5.1 and JN.1 in all the vaccinated BTI groups without RI), as well as lower than antibody response against Omicron subvariant BA.5 with titers of 2730 (IQR: 1371 to 4555) (P< 0.05 for JN.1 in all the vaccinated BTI groups without RI) and XBB with ELISA titers of 2517 (IQR: 1176 to 3750) (P<0.01 for EG.5; P<0.0001 for JN.1) (Fig. 2A, S1A).

**Figure 2.**
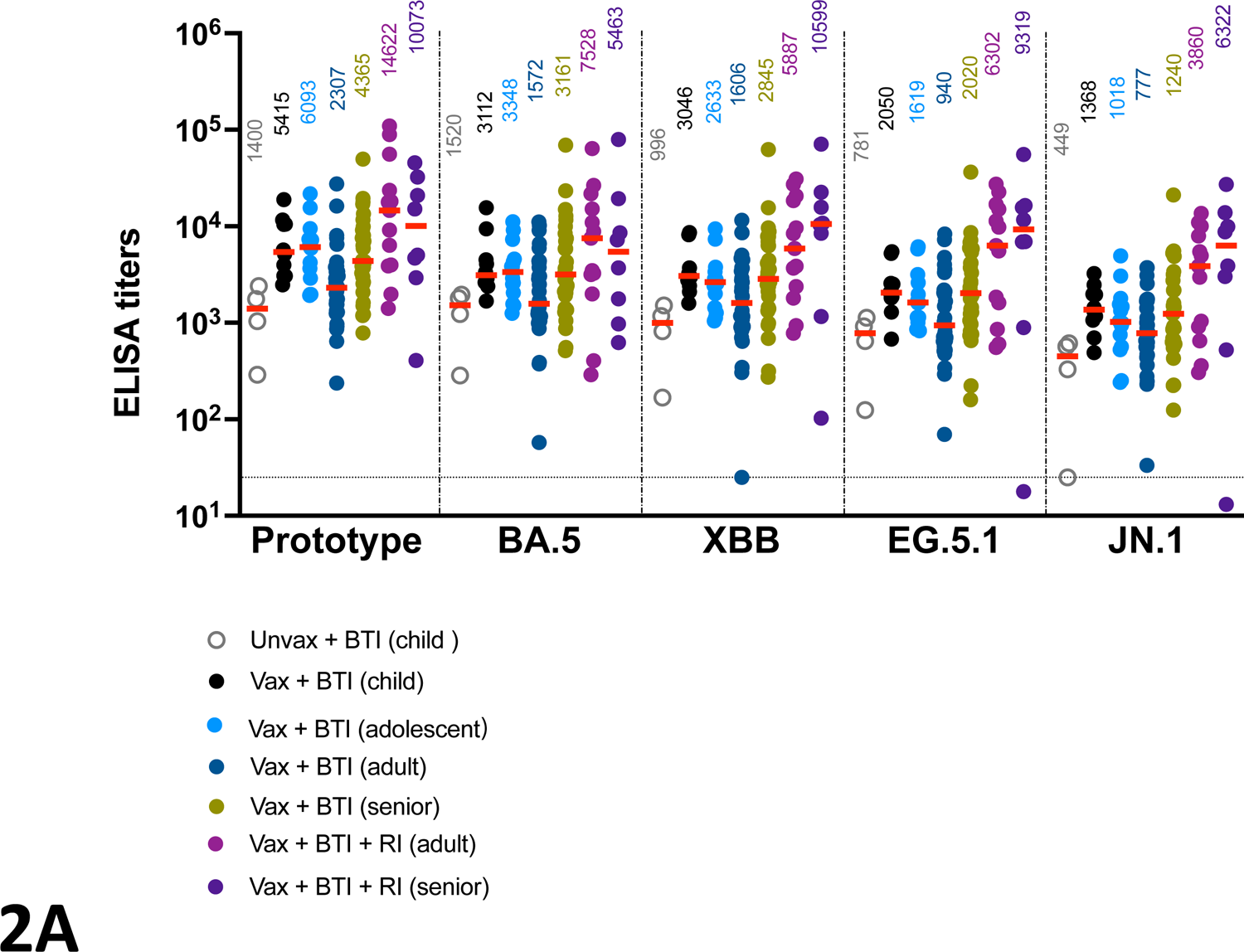
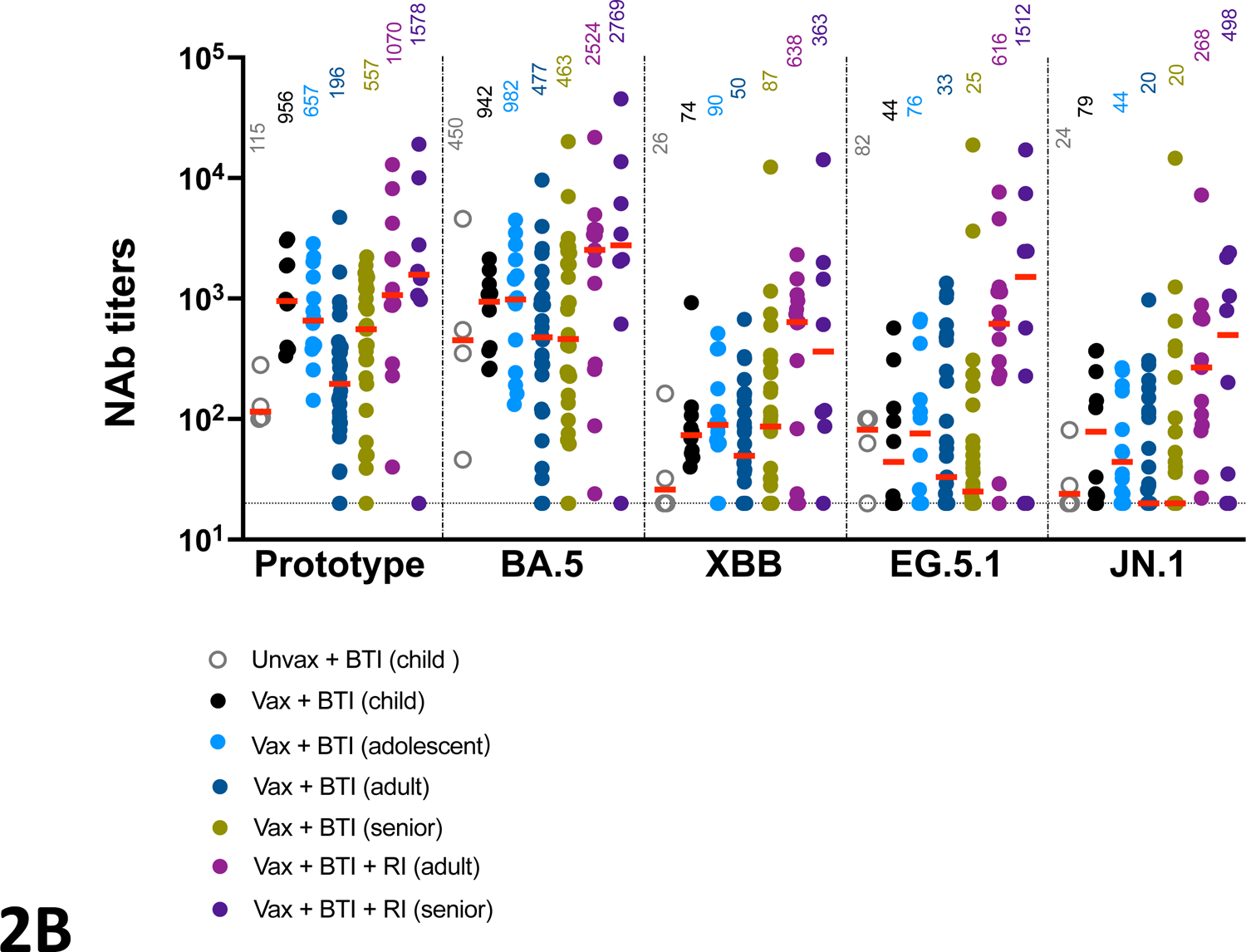
Circulating antibodies against SARS-CoV-2 following BA.5-wave breakthrough infection. Comparison of SARS-CoV-2 Spike RBD IgG titers (**A**) and SARS-CoV-2 pseduovirus neutralizing antibody titers (**B**) in response to prototype strain and Omicron lineages including BA.5, XBB, EG.5.1 and JN.1 in blood from subjects grouped as: unvaccinated children with BTI (n=4); vaccinated children with BTI (n=10); vaccinated adolescents with BTI (n=14); vaccinated adults with BTI (n=14); vaccinated adults with BTI (n=30); vaccinated seniors with BTI (n=30); vaccinated adults with BTI plus RI (n=13); vaccinated seniors with BTI plus RI (n=8). Dotted lines reflect lower limits of quantitation. Bold horizontal lines and numbers above reflect median values.

Subsequently, the neutralizing antibody (NAb) titers were assessed by a pseudovirus-based neutralization assay.^21, 22, 23^ In the vaccinated BTI groups without RI, the median NAbs against EG.5.1 and JN.1 were 33 (IQR: 20 to 129) and 24.5 (IQR: 20 to 116), respectively, which primarily exhibited above-the-threshold yet low magnitude for the two strains (Fig. 2B, S1B). Besides, NAb responses against EG.5.1 and JN.1 were observed with 12- and 16-fold lower than that against prototype with median titers of 396 (IQR: 144 to 976) (P<0.0001 for both EG.5.1 and JN.1), and with 17- and 23-fold lower compared to antibody response against Omicron BA.5 with median NAb titers of 562 (IQR: 228 to 1713) (P<0.0001 for both EG.5.1 and JN.1) (Fig. 2B,S1B).

We next evaluated the humoral immunity across groups of different ages and vaccine regimens following BA.5 BTI. At 11 months post-BTI, vaccinated children with BTI primarily maintained binding antibody response against prototype and Omicron variants, with 2∼3.8-fold higher titers than unvaccinated children with BTI (P<0.01 for all variants) (Fig. 2A). Of note, 1.7∼2.3-fold higher binding antibody titers against different variants were observed in the vaccinated children as compared with vaccinated adults and seniors by 11 months following BTI (P<0.05 for prototype) (Fig. 2A).

Consistent with binding antibody profile, vaccinated children with BTI showed ∼8.3-fold higher NAb titers against SARS-CoV-2 prototype and variants than unvaccinated children with BTI (P<0.01 for prototype) (Fig. 2B, S1B). In addition, 1.3∼4.9-fold higher NAb titers against different variants were observed in the vaccinated children as compared with that in vaccinated adults following BTI (P<0.05 for prototype), suggesting a potential superior long-term hybrid antibody response to SARS-CoV-2 in children than that in adults. Notably, the NAb response targeting recently emerging JN.1 was barely detected in seniors, with median NAb titers of 20 (IQR: 20 to 84), which exhibited 2.2∼3.9-fold lower than children with titers of 78.5 (IQR: 22.3 to 78.5) and adolescents with titers of 44 (IQR: 23 to 175.5) against JN.1 variant (Fig. 2B). Although the majority of subjects in the BTI cohort showed JN.1-specific NAb titers below 100, 8 out of 10 children and 11 out of 14 adolescents exhibited detectable level of NAb titers against JN.1, as opposed to less than half of the adults and seniors with detectable JN.1-specific NAbs. Overall, the convalescent COVID-19 cohorts generally exhibited low-level NAb response against EG.5.1 and JN.1 at 11 months post-BTI, indicating limited long-term protection against emerging Omicron variants conferred by hybrid humoral response in individuals.

### Boosting of circulating antibodies against JN.1 variant conferred by XBB/EG.5-wave reinfection

In May 2023, another wave of COVID-19 surged primarily due to the emergence of Omicron sublineages XBB and EG.5. This led to several individuals experiencing reinfection, occurring just 5 months after the initial BA.5 BTI. Significantly enhanced binding antibody responses against prototype and all the Omicron subvariants were observed in adults with BTI plus RI, as shown by 6.3-fold increase of titers against prototype (P<0.01), and 4.8∼6.7-fold elevated titers against Omicron variants (P<0.05 for variants including BA.5, XBB and EG.5). Particularly, compared with adult BTI group without RI, there was 5-fold increase of binding antibody response against fastest-growing JN.1 in adults with BTI plus RI who showed JN.1-specific NAb titer of 3860 (IQR: 779 to 8997) (P<0.05) (Fig. 2A, S1A). Similarly, binding antibody responses were also boosted in senior BTI group with RI in comparison to senior BTI group without RI, as shown by 2.3-fold increase in titers against prototype, and 1.9∼5-fold increase in titers against Omicron variants, including emerging JN.1 with boosting to the level of titer 6322 (IQR: 1148 to 12840) (Fig. 2A).

In line with the ELISA results, reinfection during the XBB/EG.5 wave boosted the NAb titers against all variants, with 5-fold increase in NAb titers against prototype (P<0.01), and 5.3∼18.6-fold increased titers against Omicron variants including BA.5, XBB and EG.5 (P<0.001 for all) (Fig. 2B, S1B). Notably, as compared with adult BTI group without RI, 13.4-fold increase of NAbs against the circulating JN.1 was observed in BTI plus RI group with the median NAb titers of 268 (IQR: 84.5 to 695) (P<0.001) (Fig. 2B, S1B). Similarly, in senior cohort the additional reinfection led to a 2.8-fold increase in titers against prototype, and 4.2∼60-fold increased titers against Omicron variants, particularly emerging JN.1 with boosting to level of titer 498 (IQR: 23.75 to 1909) (Fig. 2B). Taken together, XBB/EG.5-wave reinfection greatly augmented antibodies capable of neutralizing Omicron variants including JN.1 variant.

### Antigenic cartography

The ELISA and NAb data from all cohorts were utilized to generate antigenic maps for different groups. The plots were constructed based on the antigenic unit (AU) which accordingly represents fold change in ELISA or NAb titers.^24^ Prototype and BA.5 variant constantly cluster together with median of 0.62 (IQR: 0.4-2.1) AU for binding antibodies (Fig. 3A) and 1.6 (IQR: 0.7-2.9) AU for NAbs (Fig. 3B). JN.1 variants cluster most away from prototype, with median of 2.4 (IQR: 1.8-2.6) AU for binding antibodies and 4.1 (IQR: 2.9-5.1) AU for NAbs, suggesting higher antibody resistance compared to its predecessors.

**Figure 3.**
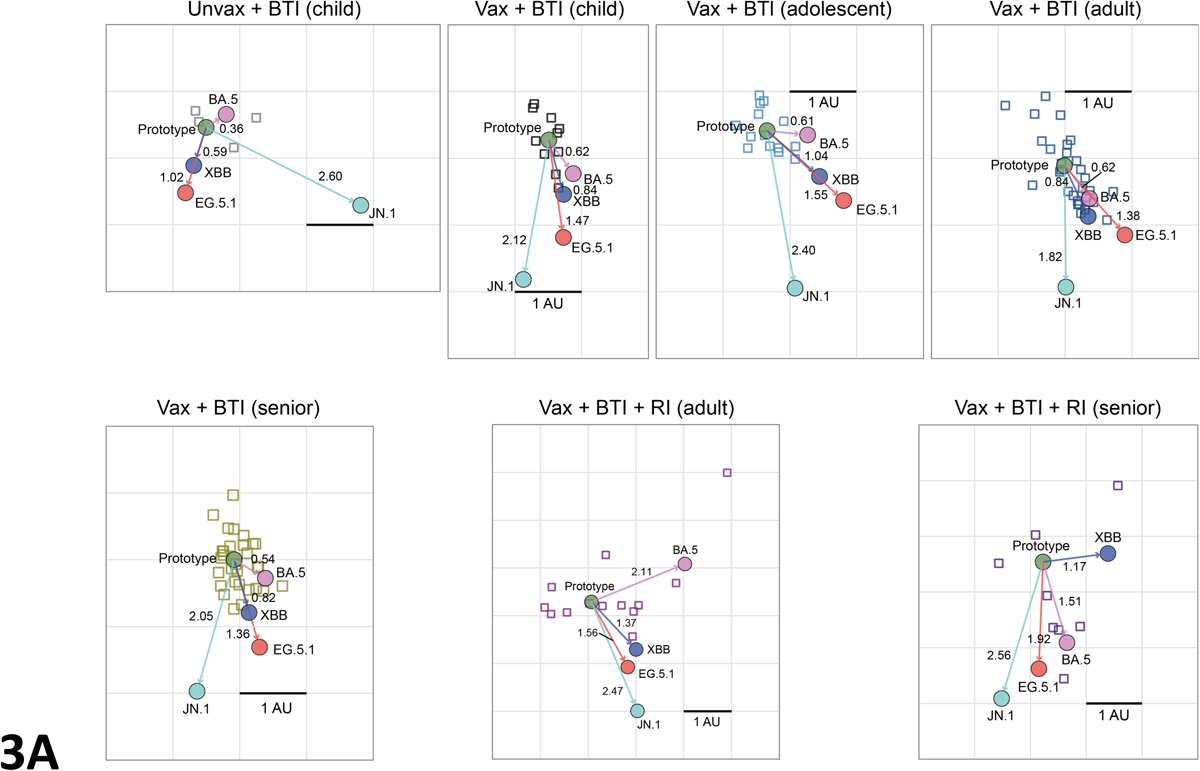
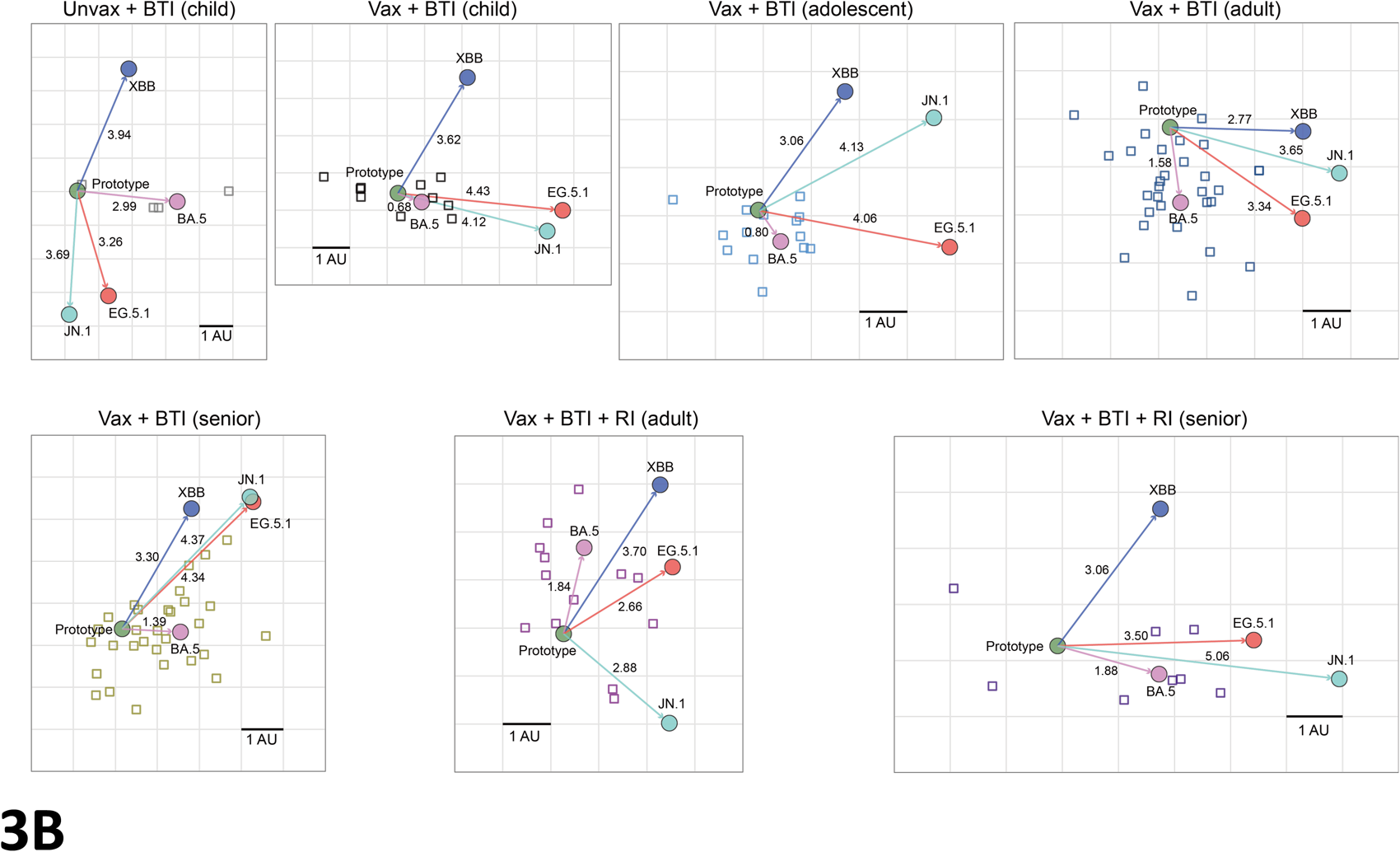
Antigenic mapping of antibody profile. The Antigenic maps were generated based on binding antibody data (**A**), and neutralizing antibody data (**B**) for all cohorts. The SARS-CoV-2 variants are shown as circles and labeled. Sera sample are shown as empty squares. Both axes represent the antigenic unit (AU) corresponding to fold change in the antibody titers.

### Enhancement of respiratory mucosal antibody responses against JN.1 through breakthrough infection

The majority of previous studies have focused on assessing humoral immunity to SARS-CoV-2 in blood samples. However, the circulating antibody profiles often do not accurately reflect the immune responses occurring in the airways. In other words, even high-level antibody response in circulation may not be necessarily corresponding to robust humoral immune protection in respiratory tract which is the primary site of entry for SARS-CoV-2.^25^ Thus, it was critical to profile respiratory mucosal antibody immune response against SARS-CoV-2, and the current understanding of vaccine and/or infection-induced mucosal immunity against Omicron lineages, particularly the emerging JN.1, in respiratory tract remains elusive.

Here, we collected BAL in parallel with blood from vaccinated subjects with BA.5-wave BTI, and examined both the circulating and respiratory mucosal neutralizing antibody activity against prototype, BA.5, XBB, EG.5 and JN.1. These vaccinated subjects with BTI all showed discernible NAb titers in blood, with significantly higher NAb titers against prototype and BA.5 than that targeting Omicron variants XBB, EG.5 and JN.1 (Prototype: P<0.01 for all; BA.5: P<0.05 for all) (Fig. S2). While no obvious difference in NAb against viral strains were noticed in BAL. Besides, paired-sample analysis indicated no correlation regarding magnitude of NAb response between blood and BAL samples. We further compared the NAb titers in BAL from vaccinated subjects with or without BTI. Individuals without BTI barely exhibited mucosal neutralizing activity against variants, while 14 out of 30 subjects with BTI showed detectable NAbs against Omicron variant XBB, EG.5 and JN.1 in BAL, with significantly higher NAb titers against BA.5 and JN.1 variant than the group without infection (P<0.05 for BA.5 and JN.1) (Fig. 4). These data suggested boosting of respiratory mucosal humoral immunity with increased potency against Omicron variants through BA.5-wave BTI.

**Figure 4.**
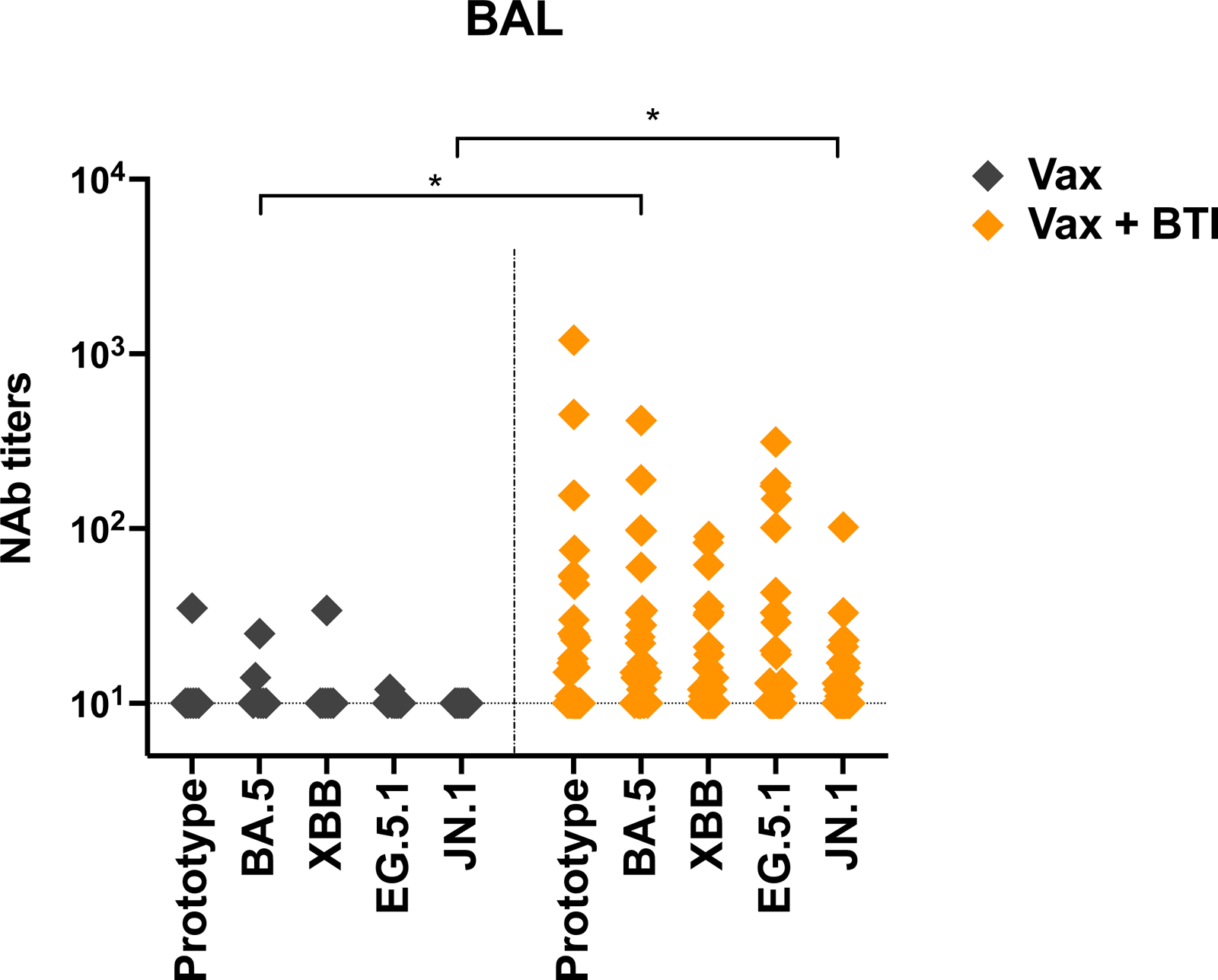
Respiratory mucosal antibody response against SARS-CoV-2 following breakthrough infection. NAb titers against prototype, BA.5, XBB, EG.5.1 and JN.1 in BAL from vaccinated subjects with BTI (orange; n=30). or without BTI (grey; n=7). Dotted lines reflect lower limits of quantitation.

### Circulating and respiratory mucosal memory B cells against prototype and JN.1 variant following breakthrough infection

While achieving sterilizing immunity against SARS-CoV-2 variants requires adequate neutralizing antibodies, protection against clinical disease can be attained through immune memory scenarios. Despite the decline in neutralizing antibody titers following SARS-CoV-2 infection, memory B cells targeting the Spike RBD not only are well-preserved in blood,^10^ but also undergo further evolution during convalescence.^26, 27, 28^ Given the durable memory B cell response against SARS-CoV-2, besides profiling the antibody response, we examined memory B cell responses against both prototype and JN.1 variant in the circulation and respiratory mucosal compartments of vaccinated individuals who experienced BA.5-wave BTI, including those with RI during the XBB/EG.5 wave.

The memory B cells responding to prototype and JN.1 variants were both detectable in blood from BTI groups with or without RI, with minimal cross-reactivity between B cells against the two strains, as indicated by the marginal presence of dual-specific B cells (Fig. 5A). The XBB/EG.5-wave RI drove the significant expansion of prototype-specific memory B cells to the level of 3.6-fold higher as compared with the group without RI (P<0.001) (Fig. 5A). Of note, there was obvious increase of B cells responding to JN.1 (P<0.05) and to both strains (P<0.05) in the RI group, with expansion to the level of 1.5-fold and 3.5-fold higher, respectively, as compared with the group without RI. Additionally, the B cells responding to prototype were markedly higher than JN.1-specific B cells (P<0.05) or dual-specific B cells (P<0.0001) in the RI group (Fig. 5A). Either the prototype-or JN.1-specific memory B cells were generally presented as resting memory (RM) pool in blood, with resting phenotype of 76% and 89% in the B cells specific to prototype and JN.1, respectively (Fig. 5B).

**Figure 5.**
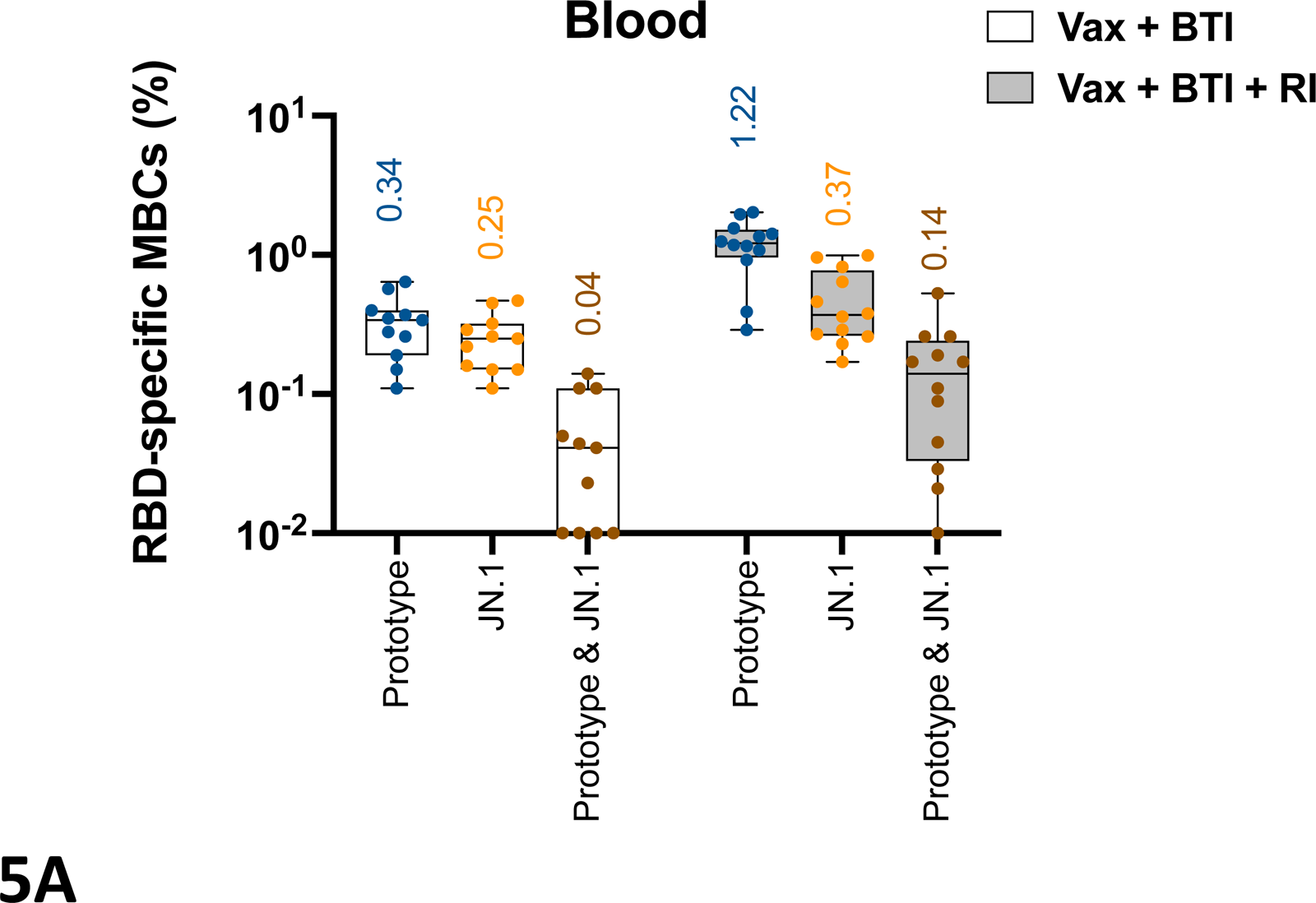
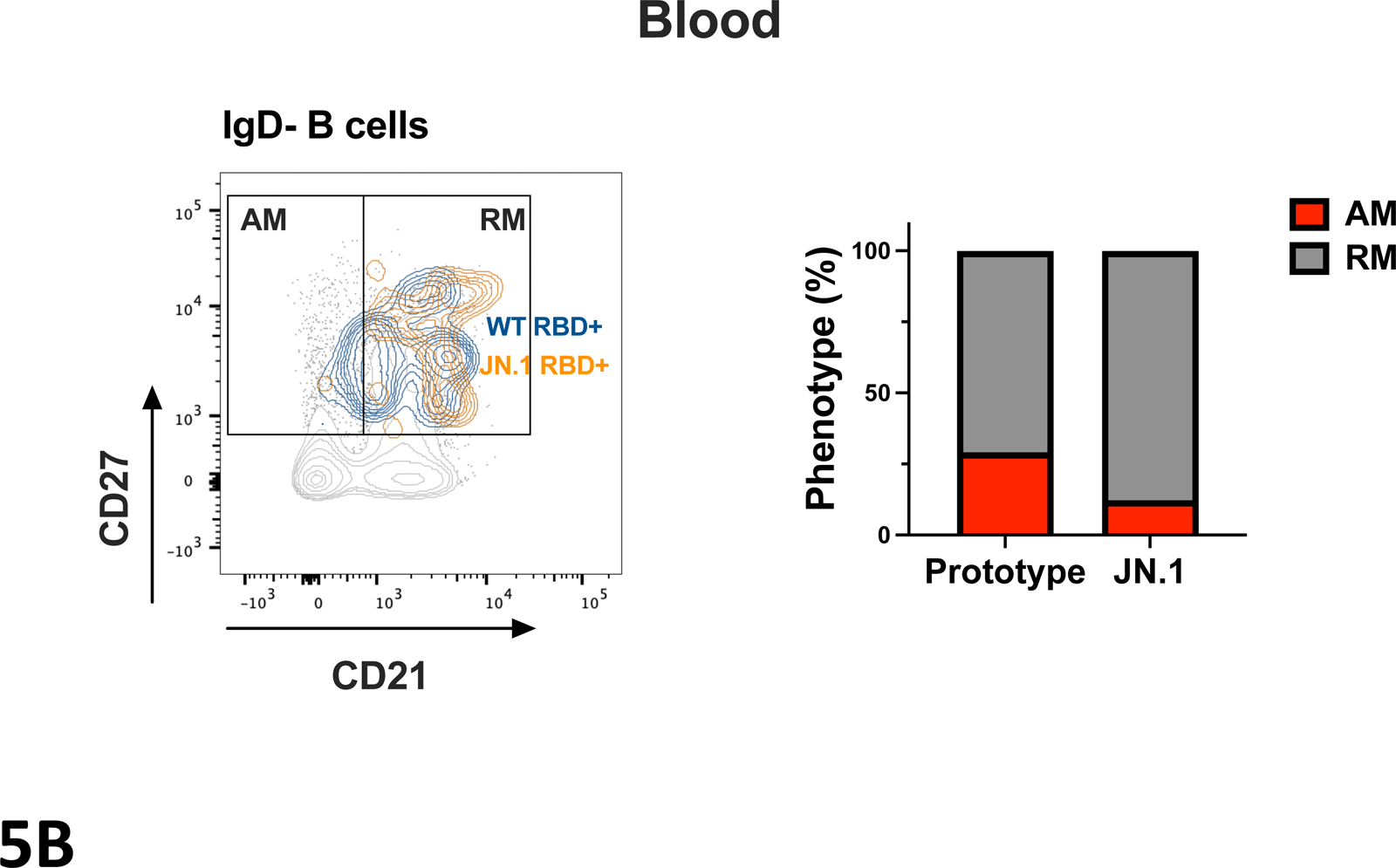
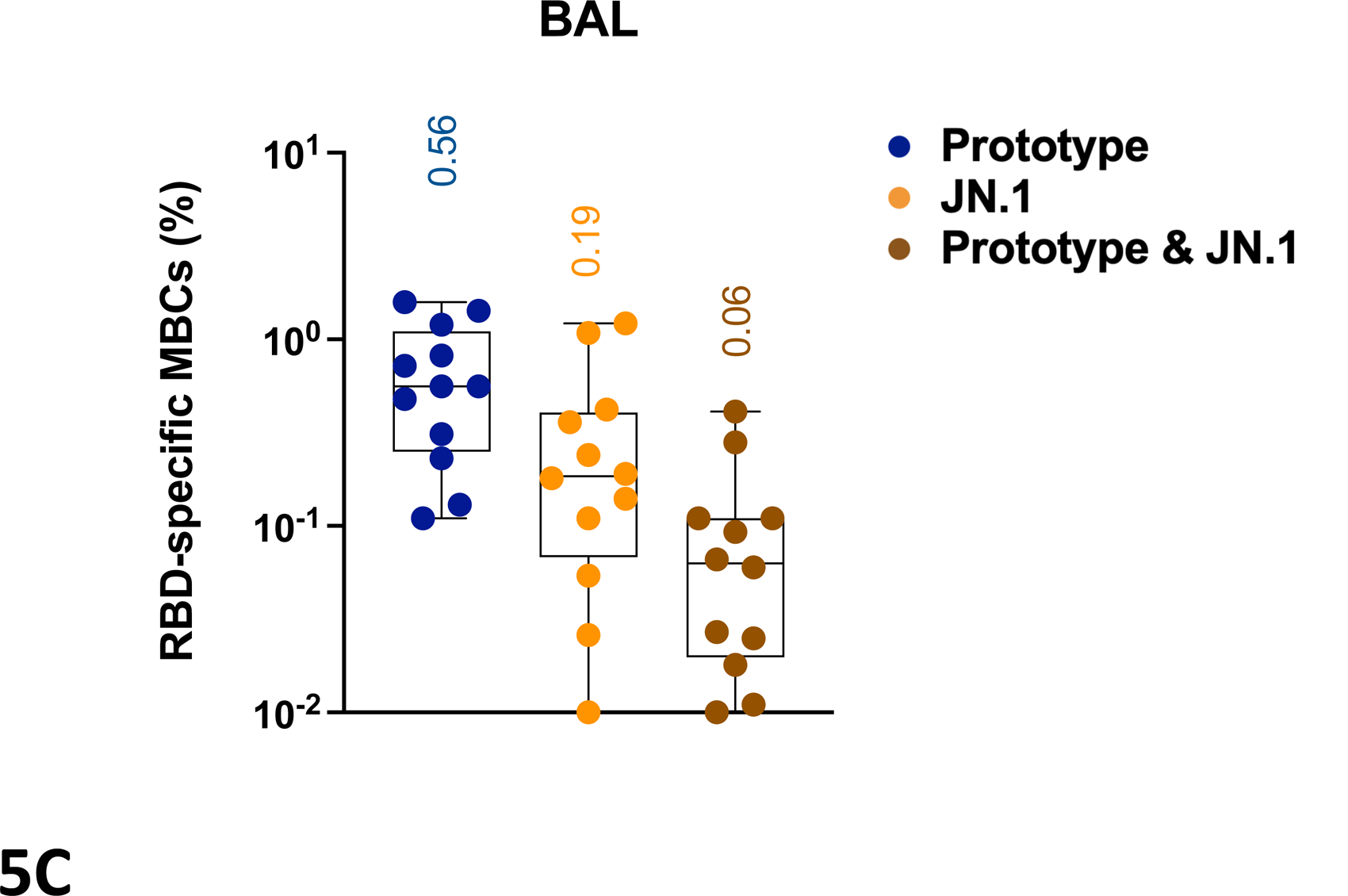
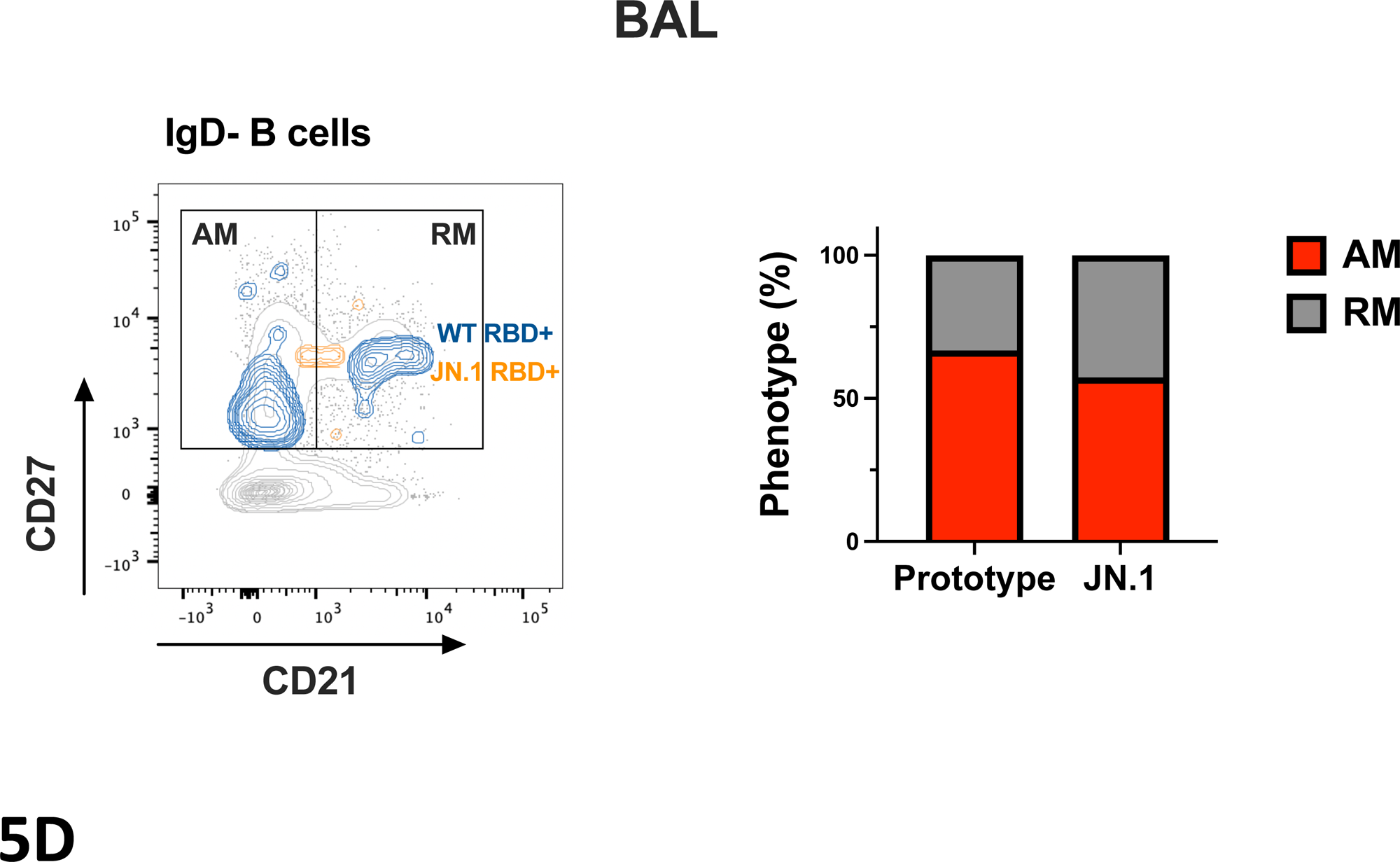
Circulating and respiratory mucosal memory B cells against SARS-CoV-2 following breakthrough infection. (**A**) Circulating memory B cells responding to prototype and JN.1 variants were measured in blood from BTI groups with RI (n=12) or without RI (n=11). (**B**) Representative flow cytometry plot show expression of CD21 and CD27 on prototype (blue) and JN.1(orange) RBD-specific memory B cells, gated on IgD-B cells in blood from subject with BTI. The phenotype of activated memory (AM; red) and resting memory (RM; grey) was indicated for circulating memory B cells responding to prototype and JN.1. (**C**) Circulating memory B cells responding to prototype and JN.1 variants were measured in BAL from BTI groups (n=12). (**D**) Representative flow cytometry plot show expression of CD21 and CD27 on prototype (blue) and JN.1(orange) RBD-specific memory B cells, gated on IgD-B cells in BAL from subject with BTI. The phenotype of AM (red) and RM (grey) was indicated for lung-resident memory B cells responding to prototype and JN.1. Box and whisker plots reflect interquartile ranges (boxes), medians (horizontal lines), and range (whiskers).

To furtherer validate the antibody-secreting function of memory B cells responding to Omicron JN.1 in BTI cohort, we assessed the JN.1-specific memory B cells by enzyme-linked immunosorbent spot (ELISpot) assay. JN.1 RBD-binding IgG-secreting memory B cells were detectable in 7 out of 11 BTI subjects with binding antibody titers above 500, which was obviously higher than that in subjects with antibody titers below 500 (P<0.05) (Fig. S3A, B). Besides, the magnitude of memory B cells secreting JN.1 RBD-binding IgG strongly correlated with binding antibody titers against JN.1 (P< 0.01, R= 0.71) (Fig. S3C).

Next, we characterized tissue-resident memory B cells responding to SARS-CoV-2 in the respiratory mucosal system by collecting BAL samples from vaccinated subjects with BA.5-wave BTI. At 11 months following BTI, memory B cells against prototype and JN.1 variant were both detectable in BAL, with higher frequency of the B cells in response to prototype than JN.1-specific B cells (P<0.05) or dual-specific B cells (P<0.0001) (Fig. 5C). Unlike the circulating memory B cells predominantly in a resting phenotype, tissue-resident memory B cells in BAL primarily exhibited activated memory (AM) phenotype with 67% in prototype-specific B cells and 57% in JN.1-specific B cells (Fig. 5D). In conclusion, memory B cells against prototype and JN.1 variant were detectable in both blood and respiratory mucosa at 11 months following BTI, with distinct memory features between the circulating and lung-resident memory B cells. XBB/EG.5-wave RI drove the significant expansion of circulating prototype-specific memory B cells, as well as increase of JN.1-specific B cells. The back-boosting of B cells responding to the ancestral viral strain highlighted the involvement of immune imprinting.

## Discussion

Due to the waning immunity and ongoing evolution of SARS-CoV-2, breakthrough infections have become common occurrences. The combinations of exposure to viral antigens during different COVID-19 waves and/or vaccination contribute to the unique immunity against SARS-CoV-2 in individuals. In the study, we undertook a thorough assessment of humoral immunity against the prototype and Omicron variants, particularly the dominant variant JN.1, in individuals with varied ages and heterologous combinations of vaccination and infection. Blood and BAL samples were collected in recent months, mostly spanning from November 2023 to January 2024. By studying the humoral immune response targeting SARS-CoV-2 in blood and respiratory mucosa, we revealed heterogeneous hybrid immunity induced by BA.5-wave breakthrough infection or XBB/EG.5 reinfection in the vaccinated cohorts.

Hybrid immunity has been reported to offer broader cross-variant neutralizing response either by inducing broad neutralizing antibodies through extensive somatic hypermutation or by targeting more conserved regions.^29, 30^ Furthermore, durable hybrid immunity was observed at 6-8 months following SARS-CoV-2 BTI, offering better protection against reinfection than immunity induced by infection or vaccination alone.^31, 32, 33^ In this study, at 11 months following BA.5-wave BTI, discernibly higher binding and neutralizing antibody responses against prototype and BA.5 variant were observed as compared to antibody responses against XBB, EG.5.1 and JN.1 across all age groups. Moreover, the NAb response against the prevalent variant JN.1 was predominantly above the threshold but remained at low level in individuals with hybrid immunity. Thus, over 11 months post-BTI, there might be potentially inadequate protection against reinfection with JN.1 which was characterized as variant of high neutralization escape.

In contrast to higher risk of clinical severity in elderly and adults, SARS-CoV-2 infection tends to manifest as mild or asymptomatic in children.^14, 15^ This suggests divergent viral and host interactions across different age groups, thereby implying the presence of heterogeneous hybrid immunity among BTI cohorts of varying ages. In this study, majority of vaccinated children and adolescents still preserved NAbs against prototype and Omicron variants including JN.1 at 11 months post-BTI, while relative lower humoral immunity occurred in adults and seniors with JN.1 NAbs mostly below the limit of detection. This finding is in line with previous studies showing that children maintained immune response against SARS-CoV-2 up to 1 year following infection.^13, 34, 35^ Our study also further indicated that the heterogeneity in hybrid antibody responses doesn’t converge into a uniform circulating antibody memory over an extended period of time post-BTI. Instead, diversity remains a prominent characteristic of humoral immunity against SARS-CoV-2 among individuals of different ages.

Recurrence of SARS-CoV-2 infection has been observed in convalescents due to the XBB/EG.5 wave with the 5-month interval following the BA.5 BTI. Notably, XBB/EG.5-wave reinfection significantly boosted the antibodies that effectively neutralize Omicron variants, including JN.1 variant, in both adults and seniors. The finding revealed the necessity of extra XBB-vaccine booster shot over 11 months following BTI, especially for the seniors who are at high risk of getting severe COVID-19 but barely maintain the NAbs against Omicron variants.

Given that SARS-CoV-2 primarily enters the host through the respiratory tract, it’s reasonable to expect that respiratory mucosal antibodies play a pivotal role as early responders to prevent viral infection. The challenge of accessing human respiratory mucosa, especially the lower airways, has posed obstacles to investigate the local respiratory immunity against SARS-CoV-2. Most studies primarily focused on humoral response against virus in human blood, which may not accurately represent the immunity in the airways. Thus, to better understand the mucosal antibody response induced by hybrid immunity over 11 months post-BTI, we profiled the NAb response against prototype and Omicron variants, including JN.1, in BAL from vaccinated subjects with or without BTI. Compared to the group without BTI that exhibited marginal Omicron-specific NAb responses, elevated mucosal NAb responses against BA.5 and JN.1 were observed in BTI cohort, particularly with 46.6% BTI individuals showing detectable JN.1-specific NAbs. These data revealed more efficient antibody protection against immune-evasive JN.1 in the lower airways conferred by BTI, which is also in line with a prior study showing that BAL samples from vaccinated individuals exhibited reduced neutralizing activity compared to convalescents.^25^

Immunological memory plays a pivotal role in providing lasting protection against pathogens. With waning antibody response along with the viral evolution over time, highly contagious Omicron variants including EG.5 and JN.1 are able to breach neutralizing humoral immunity, while memory B cells serve as a robust defense machinery against these emerging variants by establishing enduring and flexible ’antibody memory’. RBD-specific memory B cells not only maintain well in circulation for up to 8 months,^10^ but also exhibit ongoing evolution during convalescence, characterized by accumulating somatic mutations. At 6 months post-infection, the memory B cells continued clonal turnover, giving rise to expanded clones producing antibodies of elevated potency and enhanced resistance to RBD mutations. ^26, 27, 28^ Notably, at 11 months post-BTI, we observed detectable memory B cells targeting both prototype and JN.1 variant in circulation, with primarily resting phenotype. Besides, there was an increase of circulating B cells targeting JN.1 driven by XBB/EG.5-wave reinfection, along with great expansion of B cells against ancestral strain. The back-boosting with the memory B cells against original strain was attributable to immune imprinting, the downside of memory, which might limit the induction of *de novo* humoral immunity against subsequent variants. We further profiled respiratory mucosal memory B cells from BTI subjects, and validated the presence of memory B cells targeting both prototype and JN.1 in lower airways at 11 months post-BTI. Interestingly, unlike the circulating memory B cells, respiratory mucosal memory B cells predominantly displayed an activated phenotype, indicating a population primed for plasma cell differentiation. ^36^

Overall, this study has provided critical evidence elucidating the current immunity against the newly emerging JN.1 at population level. Our findings reveal the heterogeneous hybrid immunity over 11 months following breakthrough infection, and highlight the susceptibility of individuals, particularly high-risk seniors, to JN.1 breakthrough infection in the ongoing COVID-19 landscape. An additional booster might be necessary for efficient shut-down of SARS-CoV-2 transmission, and we support the official recommendation on the updated XBB-containing vaccine for the public.

## Materials and Methods

### Human cohorts

The protocol for sample collection was approved by Ethics Committee Institution at Sichuan Center for Disease Control and Prevention and West China Hospital. We recruited the cohorts of 146 subjects between May 2023 to January 2024, and the subjects were grouped by age and immunization/infection history as: unvaccinated children with breakthrough infection (BTI) (n=4; age <12); vaccinated children with BTI (n=10; age<12); vaccinated adolescents with BTI (n=14; age 12∼18); vaccinated adults with BTI (n=30; age 18∼59); vaccinated seniors with BTI (n=30; age > 60); vaccinated adults with BTI plus RI (n=13; age 18∼59); vaccinated seniors with BTI plus RI (n=8; age > 60) (Table S1). In the cohorts, all vaccinated subjects were immunized with at least one dose of inactivated vaccines of either CoronaVac (Sinovac) or BBIBP-CorV (Sinopharm) at 0.5 ml per dose, and the last dose was received between August 2021 and February 2022. All COVID-19 cases in this study were confirmed using SARS-CoV-2 PCR, antigen test kits, or serodiagnostics, and predominantly comprised mild cases. The vaccinated cohort experienced BTI during BA.5-wave from December 2022 to February 2023 in Sichuan, and samples including blood and bronchoalveolar lavage (BAL) were collected. For those subjects experiencing reinfection (RI) by XBB/EG.5-wave between May 2023 and June 2023 in Sichuan, blood samples were collected within two weeks post-symptom onset. BAL and blood samples were processed as previously described ^25, 37^.

### Enzyme-linked immunosorbent assay

The Enzyme-linked immunosorbent assay (ELISA) were conducted to measure the titers of binding antibody against the SARS-CoV-2 WT, XBB, BA.5, EG.5.1 and JN.1 (Sino Biological). ELISA plates (Corning) were coated with SARS-CoV-2 RBD protein at 1μg/mL in PBS overnight at 4 °C. The plates were then blocked using blocking buffer (PBST containing 2% BSA) at room temperature for 3 h. After washing with PBST buffer (MCE), human serum samples were serially diluted three-fold and added to the blocked plates, followed by a 1 h incubation at room temperature. Plates were washed with PBST three times. Subsequently, goat anti-human IgG antibodies (Invitrogen) were added and incubated for 1 h. Plates were washed once with PBST, and then added 100 μL per well of 3,3′,5,5′-tetramethyl biphenyl diamine (Life-iLab) and developed the plates for 10 min at room temperature in the dark. Finally, 100 μL of 1.0LM H_2_SO_4_ was added to each well, and measured the absorbance values at 450 nm on a microplate reader (Biotek).

### Production of pseudotyped lentiviral particles

The SARS-CoV-2 pseudoviruses expressing a luciferase reporter gene were generated in an approach similar to as described previously. 10 µg packaging construct psPAX2, 10 µg luciferase reporter plasmid pLenti-CMV Puro-Luc, and 5 µg spike protein expressing pcDNA3.1-SARS CoV-2 SΔCT were co-transfected into 5 x 10^6^ HEK293T cells in T-75 flask with lipofectamine 2,000 (Sigma). Six hours post-transfection, the supernatants were replaced with fresh DMEM (plus 5% FBS). The supernatants containing the pseudotype viruses were collected 48 hours post-transfection; pseudotype viruses were purified by filtration with a 0.45 µm filter.

### Lentiviral luciferase-based neutralization assay

The SARS-CoV-2 pseudoviruses neutralization assay was generated in an approach similar to that described previously^1,2^. To determine the neutralization activity of the plasma from mice, HEK293T-hACE2 cells were seeded in 96-well tissue culture plates at a density of 2 × 10^4^ cells/well overnight. Three-fold serial dilutions of heat-inactivated plasma samples were prepared and mixed with 50 µL of pseudovirues. The mixture was incubated at 37 °C for 1 h before being added to HEK293T-hACE2 cells. Moreover, 48 h after infection, cells were lysed in Firefly Luciferase Reporter Gene Assay Kits (Beyotime, RG006) according to the manufacturer’s instructions. SARS-CoV-2 neutralization titers were defined as the sample dilution at which a 50% reduction in the relative light unit (RLU) was observed relative to the average of the virus control wells.

### Antigenic cartography

The Racmacs package (https://github.com/acorg/Racmacs, version 1.2.9) was used for antigenic cartography analyses as previously described^24^. Antigenic maps were constructed based on a modified multi-scaling approach to quantify and represent the relationship between serum antibody titer data and SARS-CoV-2 variants (antigens) in a two-dimensional space. The relative positions of each SARS-CoV-2 variant and sera sample were optimized. The spacing between grid lines on the maps is one antigenic unit (AU) corresponding to fold change in antibody titers.

### B cell immunophenotyping

Fresh human BAL cells and frozen human PBMCs were stained with Aqua live/dead dye (Invitrogen) for 20min at room temperature, washed twice with 2% FBS/PBS buffer, and incubated with Fc Block (Biolegend) for 10 min at room temperature. After blocking, samples were stained with anti-CD3 (BD Biosciences, clone UCHT1, PerCP-Cy5.5, 1:200), anti-CD14 (BD Biosciences, clone M5E2, PerCP-Cy5.5, 1:50), anti-CD16 (BD Biosciences, clone 3G8, PerCP-Cy5.5, 1:100), anti-CD56 (BD Biosciences, clone B159, PerCP-Cy5.5, 1:200), anti-CD27 (BD Biosciences, clone M-T271, PE-CF594, 1:200), anti-CD38 (BD Biosciences, clone HIT2, PE-Cy7, 1:200), anti-CD21 (BD Biosciences, clone B-ly4, brilliant violet (BV) 711, 1:100), anti-CD11c (BD Biosciences, clone B-ly6, Alexa Fluor 700, 1:100), anti-CD19 (BD Biosciences, clone SJ25C1, BUV395, 1:50), anti-CD71 (BD Biosciences, clone L01.1, BUV737, 1:100), anti-IgD (BD Biosciences, clone IA6-2, APC-H7, 1:50), biotinylated SARS-CoV-2 (JN.1) spike RBD protein (Sino Biological), SARS-CoV-2 (JN.1) spike RBD protein (Sino Biological) labeled with DyLight 405, SARS-CoV-2 (2019-nCoV) spike RBD protein (Sino Biological) labeled with fluorescein isothiocyanate (FITC) and allophycocyanin (APC) for 30min at 4L. Subsequently, cells were washed twice with 2% FBS/PBS buffer, followed by incubation with BV650 streptavidin (BD Pharmingen) for 10 min at room temperature, then washed twice with 2% FBS/PBS buffer and fixed with 2% paraformaldehyde. All data were acquired on BD Fortessa flow cytometer. Subsequent analyses were performed using FlowJo software (BD Bioscience, v.10.8.1). For analyses, in singlet gate, dead cells were excluded by LIVE/DEAD Fixable Aqua Dead Cell Staining, and B cells were identified as CD19+CD3-CD14-CD16-CD56-. SARS-CoV-2 prototype-or JN.1 RBD-specific B cells were identified as double-positive for RBD proteins labeled with different fluorescent probes. The SARS-CoV-2-specific B cells were further distinguished according to CD21 and CD27 phenotype distribution: activated memory B cells (CD21-CD27+) and resting memory B cells (CD21+CD27+).

### Memory B cell ELISpot

PBMCs were cultured at 2 × 10^6^ cells/ml in R10 media (RPMI-1640 with 10% heat-inactivated FBS with 1% Penicillin/Streptomycin) supplemented with the TLR7 and TLR8 agonist imidazoquinoline resiquimod (R848, 1 μg/ml; Sigma) and 10 ng/ml recombinant human IL-2 (PeproTech) for 5-day stimulation of memory B cells. The 96-well ELISPot filter plates (Millipore) were pre-coated with 15 μg/ml capturing monoclonal anti-human IgG mAbs (Mabtech) and incubated overnight at 4°C. The plates were blocked with R10 media for minimum 30 min at room temperature. After blocking, the stimulated PBMCs were added to the plates at 0.5 million cells/well, and incubated for 28 h at 37°C. All steps following this incubation were performed at room temperature. The plates were incubated with biotinylated SARS-CoV-2 (JN.1) spike RBD protein (Sino Biological, 1 μg/ml) for 2 h and Streptavidin-AP Conjugate from Invitrogen (1.5 μg/ml) for 1 h, followed by the addition of NBT/BCIP chromogen substrate solution for 12 min. The chromogen was discarded, and the plates were washed with water and dried in a dim place for 24 h. Plates were scanned and counted on a Cellular Technologies Limited Immunospot Analyzer.

### Statistical analysis

Statistical analysis of immunologic data from the study were performed using GraphPad Prism 8.4.2 (GraphPad Software). Comparisons of immunologic data between groups were assessed using Kruskal–Wallis one-way analysis of variance. Correlations between variables were analyzed by two-sided Spearman rank-correlation tests. P values of less than 0.05 were considered significant.

## Acknowledgments

We thank Mengping Chen, Qing Yang, Jun Tang, Jiaqi Wang for generous assistance, materials and reagents.

## Author Contributions

X.H., W.M.L. designed the study. X.H., J.Y.Y., J.J.J., L.G., J.Y.L., L.L.Z., C.Y.B., Q.Z., X.L.M., S.Z., J.J.Y., C.J.Z., Y.Q.L., G.N.Z., R.J.Z. performed the immunologic assays and data analyses. D.L., Q.Q., Y.L., J.D.G., LY, J.X.W., H.H.Z., J.H.X. recruited the cohorts and processed samples. X.H., W.M.L., D.L., Z.F.W. supervised the study. X.H., J.Y.Y wrote the paper with all co-authors.

## Figure legends

**Figure S1.**
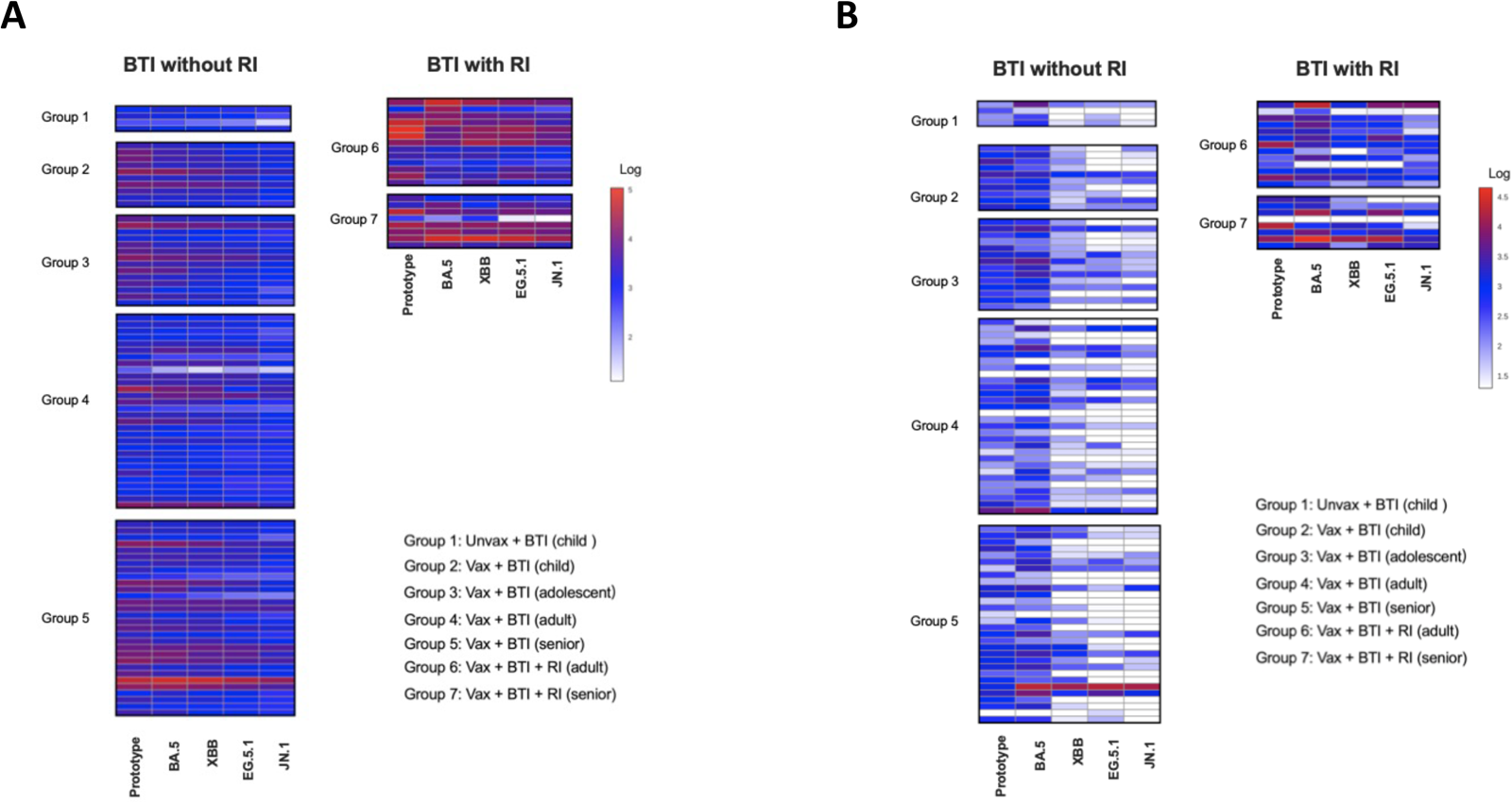
Heatmaps for SARS-Cov-2 Spike RBD-specific antibody profile following BA.5-wave breakthrough infection, related to Figure 2. Heatmaps indicated SARS-CoV-2 RBD IgG titers (**A**) and SARS-CoV-2 NAb titers (**B**) in response to prototype strain and Omicron lineages including BA.5, XBB, EG.5.1 and JN.1 in blood from BTI subjects with or without RI.

**Figure S2.**
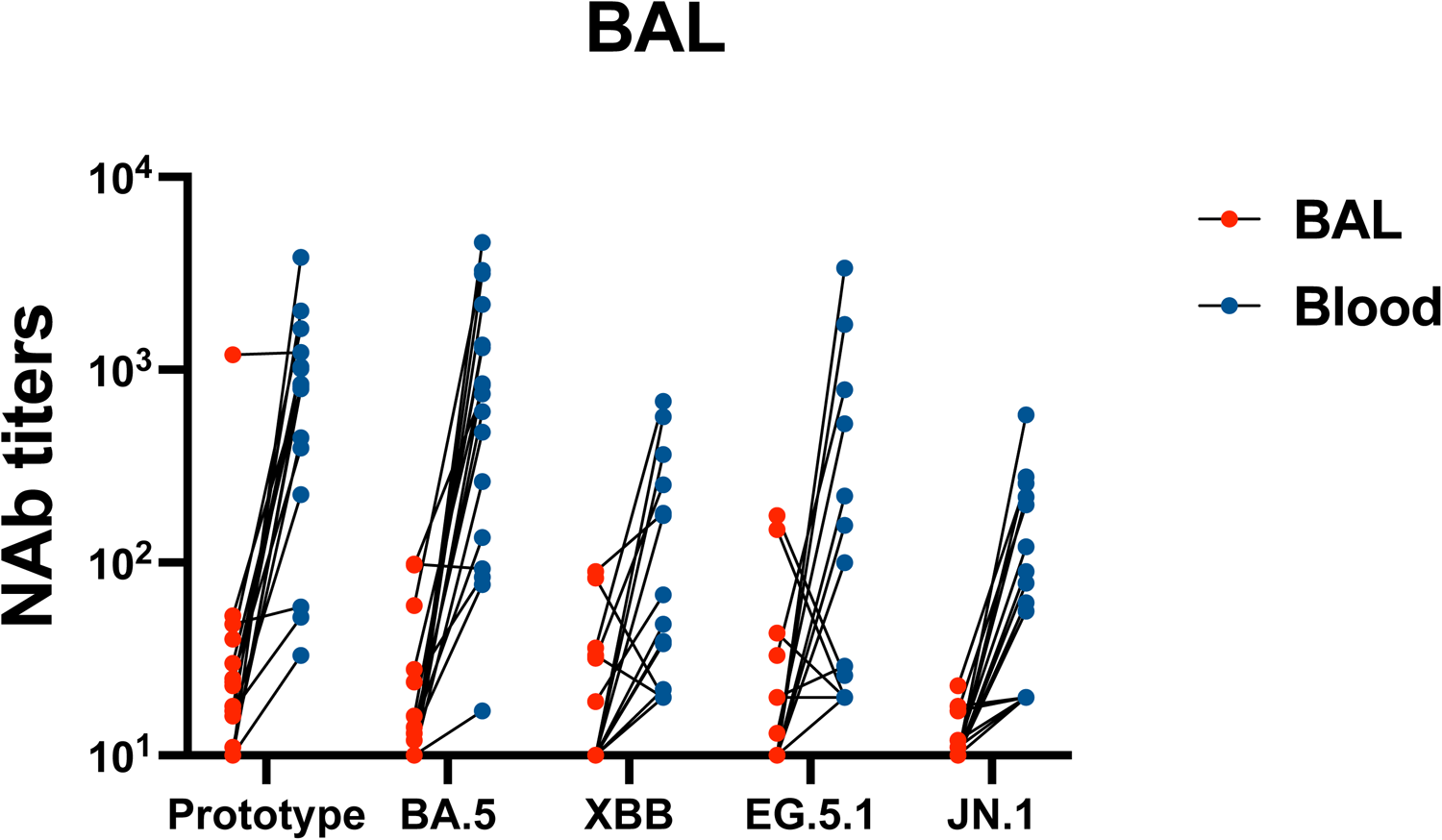
Paired-sample analysis of NAb titers in blood and BAL. Paired-sample comparison of NAb titers against prototype strain and Omicron lineages including BA.5, XBB, EG.5.1 and JN.1 was conducted for blood (blue) and BAL (red) (n=17).

**Figure S3.**
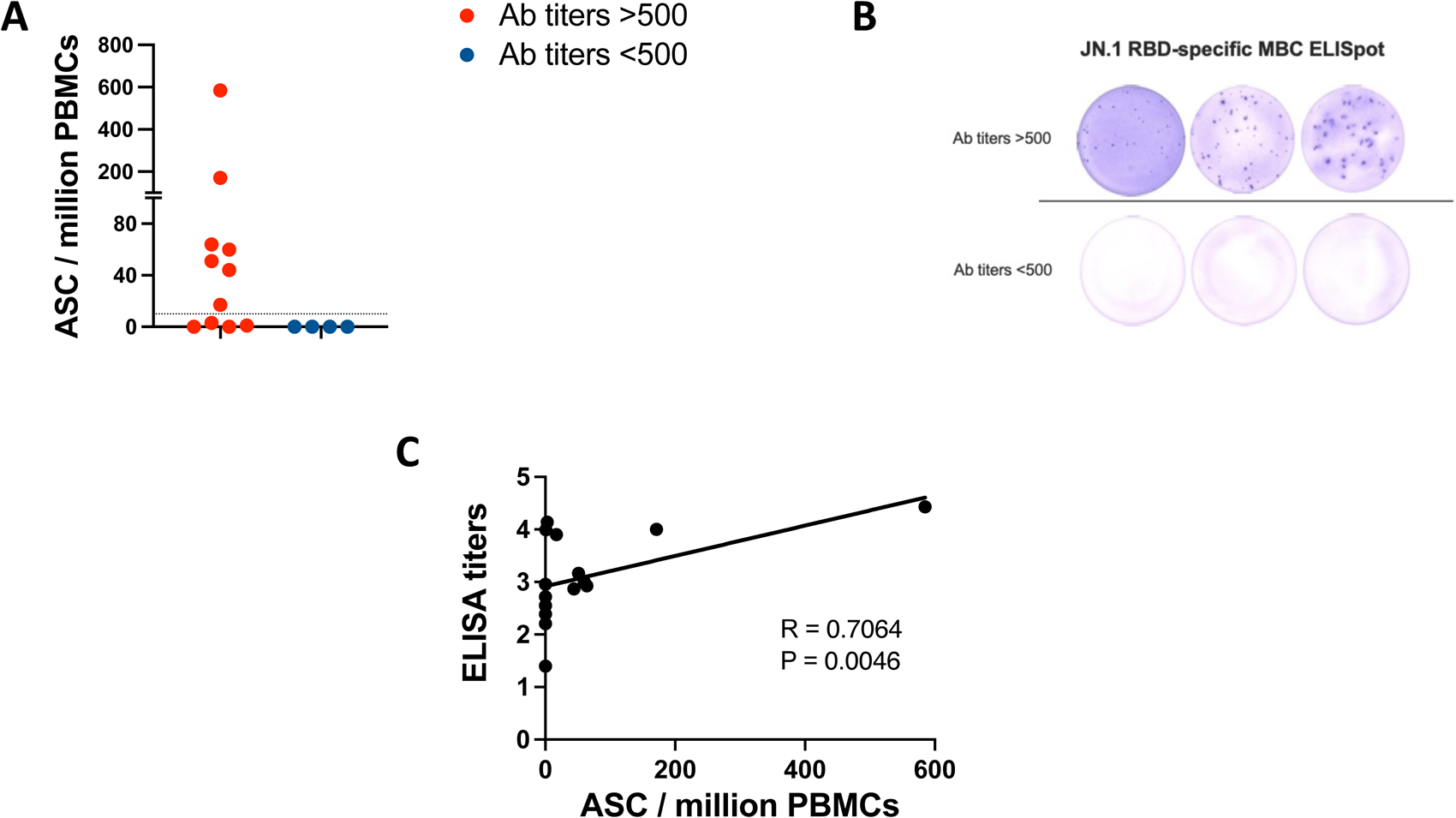
Detection of antibody-secreting function of JN.1-specific memory B cells by ELISpot. (**A**) JN.1 RBD-binding IgG-secreting memory B cells were detected in PBMCs from BTI subjects with binding antibody (Ab) titers >500 (red) or <500 (blue). Dotted line reflects lower limits of quantitation. (**B**) Representative ELISpot wells coated with JN.1 RBD and developed after plating the stimulated PBMCs from BTI subjects with binding antibody titers >500 (above) or titers <500 (below). (**C**) Correlation of JN.1 RBD-specific memory B cell ELISpot data with ELISA binding antibody titers in BTI subjects. Bold line indicates best linear fits. P and R values indicate two-sided Spearman rank-correlation tests.

Characteristics of convalescents donating blood samples
| Groups | Number of participants | Age (median, range) | Female (%) | Vaccine name | Dose |
| --- | --- | --- | --- | --- | --- |
| Group 1 (Unvax child) | 4 | 5 (5, 8) | 25 |  |  |
| Group 2 (Vax + BTI child) | 10 | 10.2 (11,4) | 40 | CoronaVac or BBIBP-CorV | 2 |
| Group 3 (Vax + BTI adolescent) | 14 | 13.5 (13.5, 4) | 42.9 | CoronaVac or BBIBP-CorV | 2 |
| Group 4 (Vax + BTI adult) | 30 | 46.2 (48, 32) | 36.7 | CoronaVac or BBIBP-CorV | 3 |
| Group 5 (Vax + BTI senior) | 30 | 67.7 (68, 19) | 43.3 | CoronaVac or BBIBP-CorV | 3 |
| Group 6 (Vax + BTI + RI adult) | 13 | 39.6 (43, 29) | 61.5 | CoronaVac or BBIBP-CorV | 1-3 |
| Group 7 (Vax + BTI + RI senior) | 8 | 76 (74.5, 33) | 50 | CoronaVac or BBIBP-CorV | 2-3 |

Characteristics of participants donating BAL samples
| Groups | Number of participants | Age (median, range) | Female (%) | Vaccine name | Dose |
| --- | --- | --- | --- | --- | --- |
| Group 1 (Vax) | 7 | 66.6 (68, 19) | 28.6 | CoronaVac or BBIBP-CorV | 2-3 |
| Group 2 (Vax + BTI) | 30 | 57.1 (60, 67) | 23.3 | CoronaVac or BBIBP-CorV | 1-3 |

## References

1. Msemburi, W., et al. The WHO estimates of excess mortality associated with the COVID-19 pandemic. Nature. 613,130–137(2023).

2. Zhang, L., et al. The significant immune escape of pseudotyped SARS-CoV-2 variant Omicron. Emerg Microbes Infect. 11,1–5(2022).

3. Qu, P., et al. Immune evasion, infectivity, and fusogenicity of SARS-CoV-2 BA.2.86 and FLip variants. Cell. 187,585–595 e586(2024).

4. Yang, S., et al. Fast evolution of SARS-CoV-2 BA.2.86 to JN.1 under heavy immune pressure. Lancet Infect Dis. 24,e70–e72(2024).

5. Wang, Q., et al. Antigenicity and receptor affinity of SARS-CoV-2 BA.2.86 spike. Nature. 624,639–644(2023).

6. Looi, M. K. Covid-19: WHO adds JN.1 as new variant of interest. BMJ. 383,2975(2023).

7. Kaku, Y., et al. Virological characteristics of the SARS-CoV-2 JN.1 variant. Lancet Infect Dis. 24,e82(2024).

8. CDC. CDC data tracker. 2024 https://covid.cdc.gov/covid-data-tracker/#variant-proportions

9. Feng, C., et al. Protective humoral and cellular immune responses to SARS-CoV-2 persist up to 1 year after recovery. Nat Commun. 12,4984(2021).

10. Dan, J. M., et al. Immunological memory to SARS-CoV-2 assessed for up to 8 months after infection. Science. 371(2021).

11. Yang, J., et al. Low levels of neutralizing antibodies against XBB Omicron subvariants after BA.5 infection. Signal Transduct Target Ther. 8,252(2023).

12. Yisimayi, A., et al. Repeated Omicron exposures override ancestral SARS-CoV-2 immune imprinting. Nature. 625,148–156(2024).

13. Jacobsen, E. M., et al. High antibody levels and reduced cellular response in children up to one year after SARS-CoV-2 infection. Nat Commun. 13,7315(2022).

14. Team, Cdc Covid-Response. Severe Outcomes Among Patients with Coronavirus Disease 2019 (COVID-19) - United States, February 12-March 16, 2020. MMWR Morb Mortal Wkly Rep. 69,343–346(2020).

15. Dhochak, N., Singhal T., Kabra S. K., Lodha R. Pathophysiology of COVID-19: Why Children Fare Better than Adults? Indian J Pediatr. 87,537–546(2020).

16. WHO. Coronavirus disease (COVID-19) Weekly Epidemiological Updates and Monthly Operational Updates. 2024 https://www.who.int/emergencies/diseases/novel-coronavirus-2019/situation-reports

17. Jeworowski, L. M., et al. Humoral immune escape by current SARS-CoV-2 variants BA.2.86 and JN.1, December 2023. Euro Surveill. 29(2024).

18. Rogers, T. F., et al. Isolation of potent SARS-CoV-2 neutralizing antibodies and protection from disease in a small animal model. Science. 369,956–963(2020).

19. Piccoli, L., et al. Mapping Neutralizing and Immunodominant Sites on the SARS-CoV-2 Spike Receptor-Binding Domain by Structure-Guided High-Resolution Serology. Cell. 183,1024–1042 e1021(2020).

20. Robbiani, D. F., et al. Convergent antibody responses to SARS-CoV-2 in convalescent individuals. Nature. 584,437–442(2020).

21. Yu, J., et al. Deletion of the SARS-CoV-2 Spike Cytoplasmic Tail Increases Infectivity in Pseudovirus Neutralization Assays. J Virol. 95(2021).

22. He, X., et al. Low-dose Ad26.COV2.S protection against SARS-CoV-2 challenge in rhesus macaques. Cell. 184,3467–3473 e3411(2021).

23. Yu, J., et al. DNA vaccine protection against SARS-CoV-2 in rhesus macaques. Science. 369,806–811(2020).

24. Smith, D. J., et al. Mapping the antigenic and genetic evolution of influenza virus. Science. 305,371–376(2004).

25. Tang, J., et al. Respiratory mucosal immunity against SARS-CoV-2 after mRNA vaccination. Sci Immunol. 7,eadd4853(2022).

26. Gaebler, C., et al. Evolution of antibody immunity to SARS-CoV-2. Nature. 591,639–644(2021).

27. He, X., et al. A homologous or variant booster vaccine after Ad26.COV2.S immunization enhances SARS-CoV-2-specific immune responses in rhesus macaques. Sci Transl Med. 14,eabm4996(2022).

28. Zhou, Z., Barrett J., He X. Immune Imprinting and Implications for COVID-19. Vaccines (Basel). 11(2023).

29. Wang, Z., et al. Naturally enhanced neutralizing breadth against SARS-CoV-2 one year after infection. Nature. 595,426–431(2021).

30. Bates, T. A., et al. Vaccination before or after SARS-CoV-2 infection leads to robust humoral response and antibodies that effectively neutralize variants. Sci Immunol. 7,eabn8014(2022).

31. Goldberg, Y., et al. Protection and Waning of Natural and Hybrid Immunity to SARS-CoV-2. N Engl J Med. 386,2201–2212(2022).

32. Abu-Raddad, L. J., et al. Association of Prior SARS-CoV-2 Infection With Risk of Breakthrough Infection Following mRNA Vaccination in Qatar. JAMA. 326,1930–1939(2021).

33. Gazit, S., et al. Severe Acute Respiratory Syndrome Coronavirus 2 (SARS-CoV-2) Naturally Acquired Immunity versus Vaccine-induced Immunity, Reinfections versus Breakthrough Infections: A Retrospective Cohort Study. Clin Infect Dis. 75,e545–e551(2022).

34. Dowell, A. C., et al. Children develop robust and sustained cross-reactive spike-specific immune responses to SARS-CoV-2 infection. Nat Immunol. 23,40–49(2022).

35. Joshi, D., et al. Infants and young children generate more durable antibody responses to SARS-CoV-2 infection than adults. iScience. 26,107967(2023).

36. Lau, D., et al. Low CD21 expression defines a population of recent germinal center graduates primed for plasma cell differentiation. Sci. Immunol. 2, eaai8153 (2017)

37. Tan CS., et al. Durability of Heterologous and Homologous COVID-19 Vaccine Boosts. JAMA Netw Open. 2022 Aug 1;5(8):e2226335.

